# siRNA biogenesis, DNA methylation and target locus silencing are distinct and separable Pol IV sub-functions differentially dependent on the largest subunit's CTD

**DOI:** 10.1101/164285

**Authors:** Jered M. Wendte, Jeremy R. Haag, Olga M. Pontes, Jasleen Singh, Sara Metcalf, Craig S. Pikaard

**Author notes:** Lead contact/corresponding author.

## Abstract

Plant nuclear multisubunit RNA polymerase IV plays a key role in the RNA-directed DNA methylation (RdDM) pathway for transcriptional silencing of transposons, viruses and specific genes by synthesizing precursors of 24 nt siRNAs that guide the process. The Pol IV largest subunit, NRPD1 is derived from the Pol II largest subunit but has a unique carboxy-terminal domain (CTD) of unknown function. We show that the NRPD1 CTD is critical for transcriptional silencing of target loci and for producing 24 nt siRNAs at high levels. However, the CTD is surprisingly dispensable for near wild-type levels of Pol IV-dependent genomic cytosine methylation. These results suggest that low levels of 24 nt siRNAs, produced at only 20-30% of wild-type levels, are sufficient for full RNA-directed DNA methylation, yet insufficient for silencing, suggesting additional roles for siRNAs beyond DNA methylation. Moreover, at a subset of target loci, neither siRNA levels nor cytosine methylation are impaired upon deletion of the CTD, yet silencing is lost. Collectively, the non-linear relationships between siRNA levels, cytosine methylation and silencing suggest the existence of additional mechanisms of silencing dependent on Pol IV transcription and mediated by the CTD, such as promoter occlusion to inhibit the activities of other polymerases.

## INTRODUCTION

Plants encode five functionally distinct nuclear multisubunit RNA polymerases: the ubiquitous Pols I, II, and III, present in all eukaryotes, and two plant-specific polymerases, Pol IV and Pol V that have specialized roles in RNA-mediated transcriptional silencing. Pols IV and V are best understood in the context of RNA-directed DNA methylation (RdDM), which is utilized by plants to silence transposons and other repeats in an RNA-guided manner (reviewed in: MATZKE AND MOSHER 2014; WENDTE AND PIKAARD 2017). In the canonical RdDM pathway, Pol IV partners with the RNA-dependent RNA polymerase, RDR2, to produce short double-stranded RNAs (LAW *et al.* 2011; HAAG *et al.* 2012; BLEVINS *et al.* 2015; ZHAI *et al.* 2015). These double stranded RNAs serve as precursors for 24 nt siRNAs, which are produced by the Dicer endonuclease, DCL3 (XIE *et al.* 2004; BLEVINS *et al.* 2015). The siRNAs are then bound by Argonaute family proteins, primarily AGO4 or AGO6, to form AGO-siRNA silencing complexes (ZILBERMAN *et al.* 2003; ZHENG *et al.* 2007). In parallel, Pol V independently transcribes the genomic regions that will be silenced, producing longer non-coding RNAs that serve to recruit AGO-siRNA complexes via complementary base-pairing interactions between the siRNAs and the Pol V transcripts (WIERZBICKI *et al.* 2008; WIERZBICKI *et al.* 2009), or the corresponding DNA (LAHMY *et al.* 2016). This allows recruitment of additional proteins, including the *de novo* cytosine methyltransferase, DRM2, and other chromatin modifiers that mediate repressive histone modifications and changes in chromatin that bring about transcriptional silencing (BOHMDORFER *et al.* 2014; PIKAARD AND MITTELSTEN SCHEID 2014; ZHONG *et al.* 2014).

Pols IV and V have unique largest subunits, which arose by the process of evolution following ancient duplication of the Pol II largest subunit gene, *NRPB1* (LUO AND HALL 2007; REAM *et al.* 2009; TUCKER *et al.* 2010; HAAG *et al.* 2014; WANG AND MA 2015). The Pol IV largest subunit, NRPD1, and the Pol V largest subunit, NRPE1, each retain the conserved domains A-H that are characteristic of all multisubunit RNA polymerase largest subunits, but each have unique carboxy-terminal domains (CTDs) suspected to contribute to their functional differences (PONTIER *et al.* 2005; HAAG AND PIKAARD 2011; HUANG *et al.* 2015). Whereas the NRPB1 CTD is comprised of 39 repeats of the heptad consensus YSPTSPS, the NRPD1 (Pol IV) CTD is comprised of a Defective Chloroplasts and Leaves (DeCL) domain of ∼100 amino acids, so named due to its similarity to a small family of chloroplast proteins implicated in pre-ribosomal RNA processing (KEDDIE *et al.* 1996; BELLAOUI *et al.* 2003). The NRPE1 (Pol V) CTD is much longer (∼700 amino acids) than either the Pol II or Pol IV CTDs and is comprised of multiple subdomains, including a linker region, a 17 amino acid-repeat region, a DeCL domain, and a glutamine-serine rich repeat region (PONTIER *et al.* 2005; HAAG AND PIKAARD 2011; HUANG *et al.* 2015; TRUJILLO *et al.* 2016). The presence of DeCL domains in both Pols IV and V, and the conservation of this domain in diverse plant lineages (HUANG *et al.* 2015) has suggested that the domain likely has one or more important functions. Indeed, the DeCL domain of NRPE1 has recently been shown to be required for production of Pol V transcripts *in vivo* and for RdDM at the vast majority of Pol V-dependent loci (WENDTE *et al.* 2017).

In this study, we tested the functional significance of the Pol IV CTD. To do so, we compared *nrpd1-3* null mutant plants with mutants transformed with transgenes that encode either full-length NRPD1 or NRPD1 missing its C-terminal DeCL domain (NRPD1-ΔCTD). We show that the truncated NRPD1-ΔCTD protein restores Pol IV-dependent DNA methylation to near wild-type levels at most loci, like full-length NRPD1, demonstrating that the DeCL domain of Pol IV is not critical for RdDM, unlike the DeCL domain of Pol V. However, siRNA levels are severely impaired when the CTD is deleted. Taken together, these findings indicate that most Pol IV-dependent siRNAs are dispensable, at least with respect to cytosine methylation. At an interesting subset of loci, wild-type levels of DNA methylation and siRNAs are restored by NRPD1-ΔCTD, yet silencing is not re-established. This suggests the existence of additional mechanisms for Pol IV-dependent silencing beyond siRNA biogenesis or RdDM.

## RESULTS

### The NRPD1 CTD is not required for Pol IV holenzyme assembly, nuclear localization, or RNA polymerase activity

To test the functional significance of the Pol IV CTD, *nrpd1-3* null mutants were transformed with transgenes encoding either full length NRPD1 or NRPD1 lacking its C-terminal DeCL domain (amino acids 1337-1453; referred to as NRPD1*-*ΔCTD) (Figure 1A). Each recombinant protein was engineered to include a carboxy-terminal Flag epitope tag that enabled immunoprecipitation using anti-Flag agarose beads. Immunoblot detection revealed that full length NRPD1 and NRPD1-ΔCTD are expressed and recovered at similar levels (Figure 1B; see also Figure 3C). Furthermore, each recombinant protein was found to co-immunoprecipitate with the Pol IV second-largest subunit, NRP(D/E)2 and with RDR2, indicating that neither holoenzyme subunit assembly nor RDR2 association require the CTD (Figure 1B).

**Figure 1.**
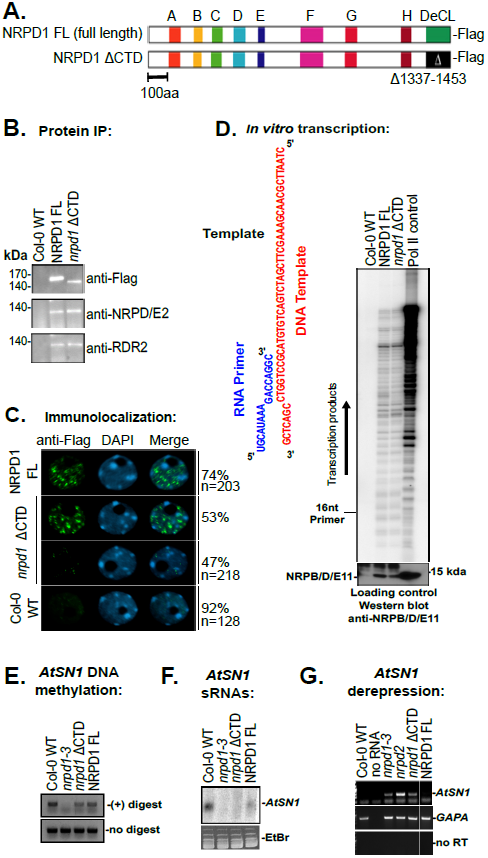
Expression and functional analysis of full length *NRPD1* and *nrpd1* ΔCTD transgenes. **A.**Schematic of the full length NRPD1 protein and NRPD1 missing the C-terminal DeCL domain. Domains A-H are conserved in all multi-subunit RNA polymerase largest-subunit genes. NRPD1 has a unique carboxy-terminal domain (CTD) characterized by a Defective Chloroplasts and Leaves (DeCL) domain similar to proteins involved in chloroplast function. Full length NRPD1 and NRPD1 with the DeCL domain deleted (deletion encompassing amino acids 1337-1453, referred to as NRPD1 ΔCTD) were transformed into *nrpd1-3* mutant plants. Both constructs incorporated a C-terminal Flag epitope. **B.**Western blot of anti-Flag immunoprecipitated full length NRPD1 or NRPD1 ΔCTD proteins. The top panel was probed using anti-Flag antibodies. The middle and bottom panels were probed using antibodies that recognize the Pol IV second-largest subunit, NRP(D/E)2, or the Pol IV interacting protein RNA DEPENDENT RNA POLYMERASE 2 (RDR2), respectively. Non-transgenic Col-0 was also subjected to immunoprecipitation as a negative control. **C.**Formaldehyde-fixed nuclei were incubated with anti-Flag antibodies to detect recombinant NRPD1 proteins. Nuclei of non-transgenic Col-0 serve as a negative control. The number of nuclei examined, and the frequency of localization patterns resembling the representative images shown, are provided to the right of the images. ***D.****In vitro* transcription assay comparing transcription activities of full length or ΔCTD NRPD1 proteins. Pol IV bound to anti-Flag resin was incubated with a DNA oligonucleotide hybridized to a RNA primer indicated in the diagram. Transcription products resulting from primer extension are labeled by virtue of incorporation of ^32^P-CTP and visualized by denaturing polyacrylamide gel electrophoresis and phosphorimaging. Immobilized flag-tagged Pol II (NRPB2-Flag) and non-transgenic Col-0 serve as positive and negative controls, respectively. Input enzyme levels were calibrated by immunoblotting using anti-NRP(B/D/E)11 to detect the 11th subunit common to Pols II, IV and V. **E.**Chop-PCR analysis of cytosine methylation at *Hae*III restriction endonuclease cut sites that undergo RdDM at the *AtSN1* locus in wild type (Col-0) plants. (+) indicates PCR conduced on genomic DNA that was digested by both enzymes and (-) indicates PCR conducted on control DNA that was subject to the same protocol but with no restriction enzyme added. Failure to detect a PCR product in the (+) digest fraction is indicative of a loss of methylation, making the DNA subject to enzymatic digestion. **F.**sRNA blot for Pol IV-dependent siRNAs derived from the *AtSN1* locus. RNA loading was controlled using ethidium bromide staining (bottom panel). **G.**RT-PCR to detect *AtSN1* transcripts, which are silenced in wild-type (Col-0). *GAPA* is a Pol II transcribed gene and serves as a loading control.

Immunolocalization using anti-Flag antibodies revealed that NRPD1-ΔCTD and full length NRPD1 are both detectable in isolated cell nuclei as punctate signals (Figure 1C), consistent with previous observations made using antibodies that recognize native NRPD1 (PONTES *et al.* 2006). However, in plants expressing the ΔCTD form of the protein, we note that the percentage of nuclei showing this punctate pattern was somewhat reduced, whereas the percentage of nuclei displaying a weak and uniform staining patter throughout the nuclei was somewhat increased (Figure 1C).

To test whether the ΔCTD deletion affects Pol IV's catalytic activity, Pol IV complexes assembled using NRPD1 or NRPD1-ΔCTD were immunoprecipitated and tested for in vitro transcriptional activity using a DNA oligonucleotide template annealed to a 16 nucleotide RNA primer (HAAG *et al.* 2012). Resulting radiolabeled RNA extension products were then resolved by denaturing gel electrophoresis and visualized by autoradiography (Figure 1D). Immunoprecipitated Pol II was tested in parallel with the Pol IV fractions, as a control. Pol II displays higher activity than Pol IV in vitro (Figure 1D), consistent with previous results (HAAG *et al.* 2012). Pol IV assembled using the NRPD1-ΔCTD subunit displayed activity indistinguishable from Pol IV assembled using full length NRPD1, indicating that the CTD is not required for Pol IV's intrinsic RNA polymerase activity (Figure 1D).

### The NRPD1 CTD is more important for siRNA levels than for cytosine methylation

For initial tests of the CTD's roles in RdDM and gene silencing, we turned to the *AtSN1* transposon family, a focus of numerous prior studies. We assays DNA methylation using Chop-PCR, a method in which genomic DNA is first incubated with a methylation-sensitive restriction endonuclease prior to attempting PCR amplification using primers that flank the endonuclease cleavage site. If the site is methylated, the DNA is not cut (chopped) and PCR amplification can occur, as is the case for DNA of wild-type Col-0 plants (Figure 1E). However, if methylation is lost, as in the *nrpd1-3* mutant, the DNA is chopped and cannot be PCR-amplified (Figure 1E). In *nrpd1* mutants expressing either full length NRPD1 or truncated NRPD1-ΔCTD, PCR products are again obtained, as in wild-type plants, indicating that Pol IV-dependent cytosine methylation has been restored by the transgenes and demonstrating that the CTD is dispensable for methylation (Figure 1E).

Given the ability of both recombinant forms of NRPD1 to rescue cytosine methylation in the *nrpd1-3* mutant background, we were surprised by RNA blot analyses that showed that full-length NRPD1 restores production of *AtSN1* siRNAs in the *nrpd1* mutant background, as expected, but NRPD1-ΔCTD does not (Figure 1F).

We next examined *AtSN1* expression, using RT-PCR, to assess the degree of transposon silencing. *AtSN1* is silenced in wild-type Col-0 plants but is derepressed in mutants defective for either the largest (*nrpd1*) or second-largest (*nrp(d/e)2*) catalytic subunits of Pol IV (Figure 1G), with the latter mutant also being defective for Pol V due to use of the same second subunit protein by Pols IV and V. We found that AtSN1 silencing is restored in *nrpd1-3* mutants that express full length NRPD1, but not in mutants expressing NRPD1-ΔCTD (Figure 1G). Taken together, these results suggest an unexpected disconnect, or non-linear relationship, among cytosine methylation, siRNA levels, and transcriptional silencing of *AtSN1* transposons.

To assess, on a genomic scale, the involvement of the CTD in Pol IV-dependent cytosine methylation and siRNA biogenesis, we conducted bisulfite sequencing in parallel with small RNA deep sequencing. Two independent biological replicates were examined for each of the genotypes tested, which were wild-type Col-0, *nrpd1-3*, or *nrpd1-3* expressing either full-length NRPD1 or NRPD1-ΔCTD (see Table S1 for complete sequencing statistics). Focusing on DNA methylation in the CHH context, which is a hallmark of RdDM, genomic regions that showed at least a 25% decrease in methylation relative to wild-type Col-0 were counted as differentially methylated regions (DMRs; see Methods). 1,916 such DMRs were identified upon comparison of Col-0 with *nrpd1-3* (Figure 2A; Table S2). Of these, 96% (1,847) had their methylation restored (to within 25% of wild type levels) by full length NRPD1 and 89% (1,701) of the DMR's had their methylation restored by NRPD1-ΔCTD (Figure 2A; Table S2), consistent with our initial Chop-PCR tests focused on *AtSN1*. Quantifying methylated CHH motif frequency as a continuous variable, displayed as the percentage of methylated CHH motifs among total CHH motifs within all Pol IV-dependent DMR regions, demonstrated that CHH methylation levels in wild-type Col-0 or *nrpd1-3* mutants expressing full length NRPD1 are indistinguishable (Figure 2B; Table S3). In *nrpd1-3* plants expressing NRPD1-ΔCTD, overall CHH methylation was restored to ∼70% of wild-type levels (Figure 2B; Table S3).

**Figure 2.**
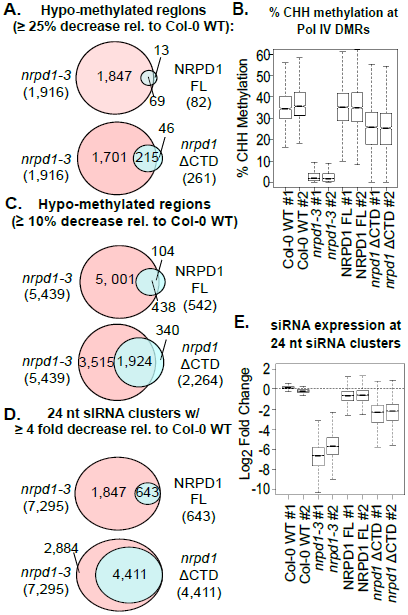
NRPD1 CTD involvement in cytosine methylation and sRNA expression. **A.**Venn diagrams comparing the number of hypo-methylated regions (regions with at least a 25% decrease in cytosine methylation relative to Col-0; see Methods) identified by whole genome bisulfite sequencing of two biologic replicates for each genotype tested. The top Venn diagram shows the number of DMRs identified in *nrpd1-3* mutants compared to the number of DMRs identified plants expressing the full length NRPD1 transgene. The bottom Venn diagram compares *nrpd1-3* to plants expressing NRPD1 ΔCTD. Genomic coordinates of hypo-methylated regions are provided in Table S2. **B.**Box plots summarizing the % methylation of all cytosines in the CHH context within differentially methylated regions (DMRs) (hypo-methylated at least 25% in *nrpd1-3* relative to Col-0; see Methods). See Table S3 for % methylation values for each DMR. **C.**Venn diagrams comparing the number of hypo-methylated regions defined using a ≥ 10% decrease in cytosine methylation as the cut-off (compare to A). See Table S2 for genomic coordinates of hypo-methylated regions. **D.**Venn diagrams comparing the number of 24 nt siRNA clusters undergoing at least 4 fold decrease in expression, relative to the average expression in Col-0. The top diagram shows the number of these regions identified in *nrpd1-3* mutants compared to plants expressing the full length NRPD1 transgene and **A.**the bottom panel compares *nrpd1-3* to plants expressing NRPD1 ΔCTD. See Table S4 for genomic coordinates of siRNA clusters. **E.**Box plots summarizing the log_2_ fold change in siRNA expression relative to the average expression in Col-0 WT across all 24 nt siRNA clusters for each replicate of each genotype tested. See Table S4 for fold change values at all regions.

Re-examination of CHH methylation using more relaxed criteria, in which a decrease in CHH methylation of only 10% qualifies as a DMR, 5,439 DMRs were identified in *nrpd1-3* mutants compared to wild type (Figure 2C; Table S2). Full-length NRPD1 rescued methylation at ∼90% of these DMRs, but 542 regions remained at least 10% hypo-methylated (Figure 2C; Table S2). The *NRPD1*-Δ*CTD* transgene also rescued methylation at the majority of loci (∼60%), but 2,264 regions remained at least 10 % hypomethylated (Figure 2A; Table S2). The results of the stringency tests suggest that at those loci where the CTD influences CHH methylation levels, methylation losses are partial and quantitative rather than absolute, unlike the effects of a null mutant.

Analysis of the RNA-seq data identified 7,295 genomic loci where 24 nt siRNA clusters occur and where siRNA read counts decrease at least 4-fold in *nrpd1-3* mutants relative to wild-type Col-0 (Figure 2D, Table S4). Expression of full-length NRPD1 restored the siRNA levels at ∼91% of these clusters, but 643 clusters continued to have siRNA levels reduced at least 4-fold relative to wild-type (Figure 2D; Table S4). In contrast, upon expression of NRPD1-ΔCTD, siRNA levels for 60% of the loci (4,411/7,295) remained at least 4-fold lower than in wild type (Figure 2D; Table S4). Quantification of siRNA read counts, summed for all 24 nt siRNA clusters, revealed that in *nrpd1-3* mutants, small RNA levels drop to ∼1% of wild-type (log_2_ ratios of −6.6 and −5.7, relative to wild-type, for the two replicates; Figure 2E; Table S4). Expression of full-length NRPD1 substantially restored siRNA levels at these loci, to ∼70% of wild-type (log_2_ ratios of −0.7 to −0.6 for the two replicates). However, expression of NRPD1-ΔCTD also restored siRNA levels to only to ∼20% of wild-type (log_2_ values of −2.3 and −2.2 for the two replicates; see Figure 2E; Table S4).

### The majority of Pol IV-dependent RNAs are dispensable for RNA directed DNA methylation

To more closely examine the relationship between siRNA and methylation levels, we identified the 500 loci at which the % CHH methylation is most similar in wild-type plants and in *nrpd1-3* mutants expressing NRPD1-ΔCTD. At these loci, methylation is severely depleted in *nrpd1-3* mutants yet is completely rescued by either full-length NRPD1 or NRPD1-ΔCTD (Figure 3A). However, siRNA levels at these 500 loci are not fully restored by NRPD1-ΔCTD; instead, their levels remain ∼3-fold lower than in wild-type (Figure 3A; Table S5). Taken together, the methylation and siRNA data suggest that two-thirds of the siRNAs produced in wild-type cells may be dispensable, without impacting the maximal level of methylation achievable by RdDM at the corresponding loci.

**Figure 3.**
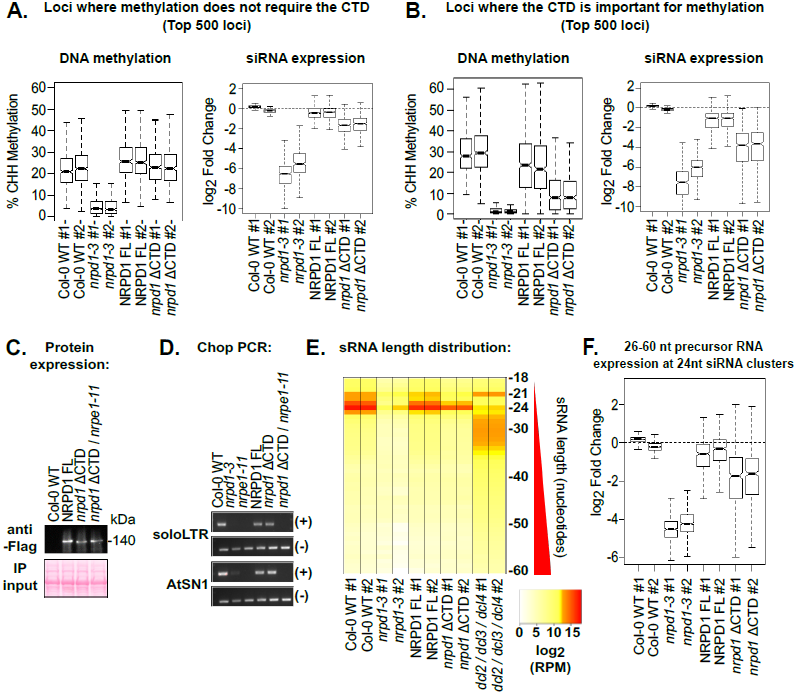
Evidence that most Pol IV-dependent siRNAs are not required for RdDM. Loci that undergo Pol IV-dependent DNA methylation were ranked based on the degree to which NRPD1 ΔCTD and WT Col-0 differ. The top 500 loci, where methylation levels were fully restored to the highest extent were analyzed in **A** and the 500 loci where methylation was least restored are shown in **B**. In A and B, the left panel shows box plots summarizing % methylation and the right panel shows summaries of log_2_ fold change in siRNA expression at the 500 loci for the two replicates for each genotype. See Table S5 and S6 for sRNA fold change values for individual loci included in the anaylses. **C.**Immunoblot affinity captured NRPD1 proteins in different genetic backgrounds. Equal input levels were calibrated with Ponseau S staining of nitrocellulose membranes. **D.**Chop-PCR analysis of cytosine methylation at *Alu*I or *Hae*III restriction endonuclease cut sites that undergo RdDM in wild type (Col-0) plants. (+) indicates PCR conduced on genomic DNA that was digested by both enzymes and (-) indicates PCR conducted on control DNA that was subject to the same protocol but with no restriction enzyme added. Failure to detect a PCR product in the (+) digest fraction is indicative of a loss of methylation, making the DNA subject to enzymatic digestion. **E.**Heatmap depicting the distribution of sRNA size classes within 24 nt siRNA clusters. Values depicted are the log_2_ of reads per million mapped for each size class. See Table S7 for RPM values of each size class. **F.**Box plot showing the log_2_ fold change in levels of RNAs sized 26-60 nt in regions corresponding to 24 nt siRNA clusters in wild type plants. Regions in which a minimum coverage of 10 reads for combined **C.**precursor size classes in each Col-0 replicate were included in the analysis. See Table S8 for regions and fold change values.

We next examined the 500 loci where methylation is least rescued, relative to wild-type, by NRPD1-ΔCTD. At these loci, siRNAs levels in the NRPD1-ΔCTD plants are ∼6-fold lower than WT (Figure 3B, Table S6). Collectively, our observations suggest that a 3-fold reduction in siRNA levels can be tolerated without detectable loss of methylation, but a 6-fold reduction in siRNAs correlates with reduced levels of cytosine methylation, suggesting the possibility of a threshold level somewhere in between.

We considered the possibility that the CTD may enable Pol IV function via an alternative RdDM pathway. As a test of this hypothesis, we crossed *nrpd1* plants expressing full-length NRPD1 or NRPD1-ΔCTD with an *nrpe1-11* mutant defective for Pol V. As expected, the expression level of the NRPD1-ΔCTD protein was unaffected by the *nrpe1* mutation (Figure 3C). Assessment of DNA methylation using Chop-PCR assays revealed that methylation was lost at *AtSN1* and *soloLTR* loci in both *nrpd1-3* and *nrpe1-11* mutants and was restored in *nrpd1* mutants that express full length or ΔCTD versions of NRPD1 (Figure 3D). However, methylation is not restored if *nrpe1* is also mutant, indicating that NRPD1 transgene-dependent restoration of methylation is still Pol V-dependent, consistent with methylation via the known RdDM pathway.

We next tested whether the reduced siRNA levels in *nrpd1* ΔCTD could be attributed to impaired processing of Pol IV and RDR2-dependent siRNA precursors (P4R2 RNAs) into siRNAs. P4R2 RNAs are mostly ∼30-50 nt, double stranded RNAs (BLEVINS *et al.* 2015; ZHAI *et al.* 2015). Full length P4R2 RNAs are efficiently processed by Dicers into siRNAs, and are thus present at low levels in wild type plants (BLEVINS *et al.* 2015). However, when siRNA processing is impaired, as in Dicer mutants, the longer P4R2 RNAs accumulate (BLEVINS *et al.* 2015; LI *et al.* 2015; ZHAI *et al.* 2015). We examined the size distribution of sRNAs in our libraries, as well as sRNAs sequenced from *dcl2 / dcl3 / dcl4* triple mutants (BLEVINS *et al.* 2015). In wild-type plants, the siRNA size distribution shows a strong peak at 24 nt and low levels of higher molecular weight size classes (Figure 3E; Table S7). Pol IV-dependent siRNAs are virtually eliminated in *nrpd1-3* mutants, but are restored upon expression of full-length NRPD1 or NRPD1-ΔCTD (Figure 3E; Table S7). This is in contrast to *dcl2 / dcl3 / dcl4* triple mutants in which 24 nt siRNAs are lost and P4R2 RNAs longer than 24 nt accumulate, with a peak at ∼30 nt (Figure 3E; Table S7) (BLEVINS *et al.* 2015). Collectively, the restoration of 24 nt siRNAs, without evidence for an increase in precursors, indicates that dicing is not disrupted in plants expressing NRPD1-ΔCTD. This, in turn, suggests that the low levels of siRNAs in NRPD1-ΔCTD plants results from reduced precursor RNA synthesis. Consistent with this hypothesis, examination of putative P4R2 RNAs in the size range of 26-60 nt, which can be detected at low levels at some loci in wild-type plants, revealed that their levels are reduced in plants expressing NRPD1*-*ΔCTD to an extent that is proportional to the reduction in siRNA abundance (Figure 3F; Table S8).

### Pol IV is required for transcriptional silencing independent of DNA methylation

In *nrpd1* mutants, ∼50 transposable element families or protein coding genes are derepressed, including *AtSN1* transposons (BLEVINS *et al.* 2014). We investigated the expression status of 15 of these loci to assess NRPD1 CTD involvement in their transcriptional silencing. In wild-type plants, transcripts derived from these 15 loci are either not detectable or present at low levels (Figure 4; Figure 5). By contrast, all 15 are expressed at high levels in *nrpd1-3* mutants, as detected by RT-PCR, indicating the loss of silencing (Figure 4; Figure 5). Expressing full-length NRPD1 in the *nrpd1-3* mutant background restores silencing to wild type levels for all loci tested (Figure 4; Figure 5). Interestingly, expression of NRPD1-ΔCTD failed to restore silencing at 7 of the 15 loci tested (*AtSN1, SDC, AT3TE63155, AT5TE47755, AT3TE47840, ERT14*, and *soloLTR;* Figure 4A-G), yet did restore silencing at 8 of the loci (*ERT7, ERT9, AT3G30844, AT1TE75910, AT2TE07550, AT2TE82000, AT3TE37570*, and *AT4TE27915*; Figure 5A-H). We considered the possibility that loci where NRPD1-ΔCTD was unable to restore silencing might correspond to loci where wild-type levels of DNA methylation are also not re-established. However, this is not the case; methylation levels were surprisingly equivalent, or nearly equivalent (less than a 10% decrease) to wild-type (Figure 4). Collectively, these suggests indicate that the CTD is needed for silencing in a manner that is independent of DNA methylation.

**Figure 4.**
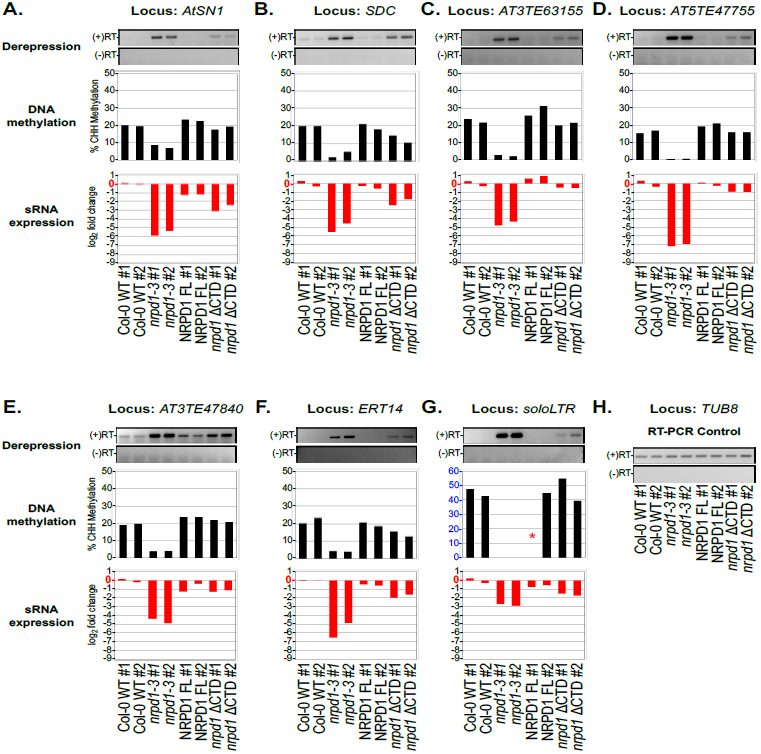
DNA methylation and sRNA expression profiles of genes that require the Pol IV CTD for transcriptional silencing. RNA expression at loci that require Pol IV for transcriptional silencing were analyzed by RT-PCR, in parallel with analysis of their CHH methylation and siRNA levels. **A.** *AtSN1* (*AT3TE63860*); **B.** *SDC* (*AT2G17690*); **C.** *AT3TE63155*; **D.** *AT5TE47755*; **E.** *AT3TE47840*; **F.** *ERT14 (AT2G01422)*; **G.** *soloLTR (AT5TE35950)*; **H.** *TUB8*. For each locus, the top panel shows RT-PCR to detect derepression visualized on agarose gels, the middle panel shows % CHH methylation, and the bottom panel shows the log_2_ fold change in siRNA expression relative to the average expression of both Col-0 WT replicates. *TUB8* (tubulin) is a Pol II transcribed gene that is not subject to RdDM and serves as an RNA input control for RT-PCRs. There was not adequate bisulfite sequencing coverage in the NRPD1 FL replicate #1 to assess % methylation for this line at the *soloLTR* locus (indicated by *). Note also that the scale for the % methylation histogram of *soloLTR* is different than the other loci to account for higher levels of methylation at this site.

**Figure 5.**
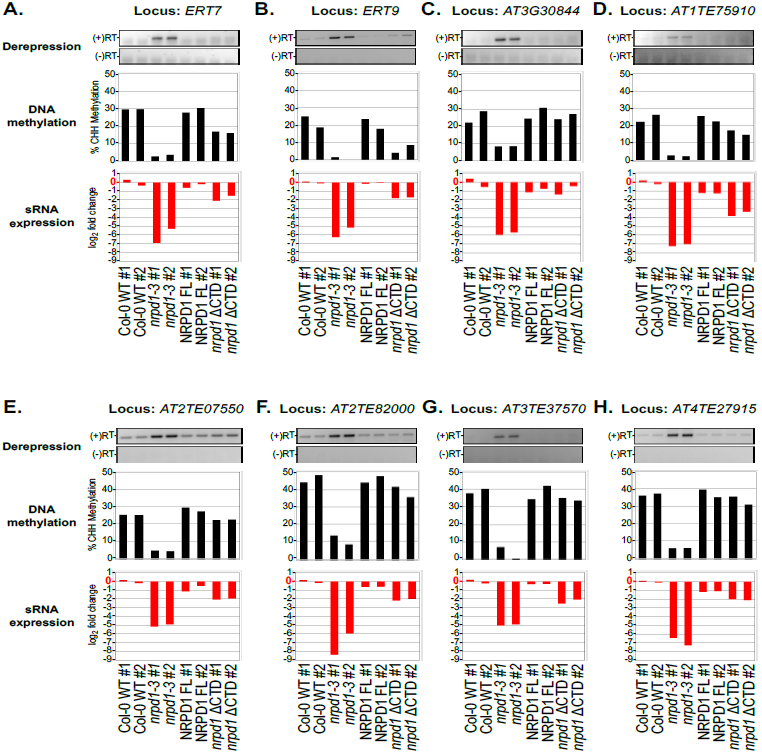
DNA methylation and sRNA expression profiles of genes that require Pol IV, but not the Pol IV CTD for transcriptional silencing. **A.** *ERT7* (*AT3G28899*); **B.** *ERT9* (*AT5G24240*); **C.** *AT3G30844*; **D.** *AT1TE75910*; **E.** *AT2TE07550*; **F.** *AT2TE82000*; **G.** *AT3TE37570*; **H.** *AT4TE27915*. For each locus, the top panel shows results of RT-PCR to detect derepression visualized on agarose gels, the middle panel shows % methylation, and the bottom panel shows the log_2_ fold change in siRNA expression relative to the average expression of both Col-0 WT replicates. The input control for the RT-PCR reactions is shown in Figure 4H.

We next asked whether there is a discernable relationship between silencing and siRNA levels at the 15 loci chosen for investigation. Most loci that failed to be silenced in NRPD1-ΔCTD plants showed a decrease in siRNA levels compared to wild type, but there were exceptions (Figure 4). For instance, at *AT3TE63155*, siRNA levels were nearly fully restored to wild-type levels (Figure 4C, 4D). Furthermore, similar, or even more severe, losses of siRNAs occur at loci where silencing is nonetheless restored by NRPD1-ΔCTD (Figure 5; see especially *AT1TE75910*, Figure 5D).

We considered the possibility that single-nucleotide, position-specific changes in small RNA or DNA methylation frequency might go unnoticed in sliding-window analyses that only count RNAs or methylcytosines within a given interval. However, displaying sRNAs and methylcytosine data using JBrowse (SKINNER *et al.* 2009) revealed no evidence that this is the case (see Figure S1 for JBrowse images corresponding to the loci examined in Figure 4 and see Figure S2 for loci examined in Figure 5).

## DISCUSSION

Our investigations into the functional significance of the Pol IV largest subunit's CTD unexpectedly revealed that siRNA levels, DNA methylation levels, and transcriptional repression are not correlated with one another in a simple, proportional manner. The CTD is most needed for production of high levels of siRNAs, such that deletion of the CTD results in overall siRNA levels that are only ∼20-30% of wild-type. Remarkably, this substantial reduction in siRNA abundance has a relatively modest effect on RNA-directed cytosine methylation. This suggests that siRNAs are produced in excess of the number needed to achieve the maximal methylation level achievable by RdDM at target loci. This revelation may have bearing on recent studies that showed that residual methylation can occur in dicer mutants defective for the production of siRNAs (STROUD *et al.* 2013; YANG *et al.* 2016; YE *et al.* 2016). These observations have been interpreted in multiple ways, with one hypothesis being that Pol IV and RDR2-dependent siRNA precursors (P4R2 RNAs) may be able to guide methylation without the need for dicing into siRNAs (YANG *et al.* 2016). Another study found that in dicer mutants, AGO4 is nonetheless loaded with small RNAs that are the same size as diced siRNAs, leading to the hypothesis that these small RNAs are generated by exonuclease trimming of P4R2 RNA precursors (YE *et al.* 2016). We propose a third, relatively simple explanation, which is that short P4R2 transcripts, falling within the bell-shaped normal size distribution of transcript sizes, whose peak is at 30 nt, just happen to be the right size to be incorporated into AGO4 without any need for dicing. Our finding in this study that siRNAs produced at low levels may direct disproportionately high levels of RdDM suggests that low levels of small RNAs have higher potency than previously suspected.

At many loci at which the absence of the Pol IV CTD causes low levels of siRNA production, yet full methylation, silencing still does not occur, suggesting that high levels of siRNAs, rather than high levels of cytosine methylation, are needed for silencing. This, in turn, suggests that siRNAs may have functions separate from, or in addition to, the guidance of DNA methylation, such as the guidance of chromatin modifying enzymes responsible for establishing repressive histone modifications, as occurs in those eukaryotes that do not methylate their genomes (HOLOCH AND MOAZED 2015; MARTIENSSEN AND MOAZED 2015). However, we also identified loci at which essentially wild-type levels of siRNAs and wild-type levels of cytosine methylation are both detected, and yet silencing does not occur in the absence of the CTD, suggesting that the CTD mediates additional mechanisms of Pol IV-dependent silencing. One possibility is that the very process of Pol IV transcription is inhibitory to transcription of genes or transposons, possibly by preventing the stable association of transcription factors needed for recruitment of RNA polymerases II or III. This latter hypothesis was put forward previously based on studies in maize that showed that LTR retrotransposons that are subjected to RdDM and whose silencing depends on Pol IV are not similarly dependent on other proteins of the RNA-directed DNA methylation pathway, such as the Snf2-like protein, RMR1, or the maize ortholog of RDR2, MOP1 (HALE *et al.* 2009). In *rmr1* or *mop1* mutants, siRNAs and cytosine methylation levels are reduced, similar to Pol IV mutants. However, transcriptional derepression of the transposons is only detected in the Pol IV mutants and not in the *rmr1* or *mop1* mutants, supporting the hypothesis that Pol IV mediates silencing independent of the RdDM process (HALE *et al.* 2009). Nuclear run-on studies in both maize and Arabidopsis subsequently revealed that in Pol IV mutants, an increase in Pol II occupancy is detected in the region just downstream of the polyadenylation sites of protein-coding genes, suggesting that Pol II termination and polymerase release is somehow induced by Pol IV activity (ERHARD *et al.* 2015; MCKINLAY *et al.* 2017). Moreover, no correlation with siRNAs or DNA methylation was found in these 3' genic regions, suggesting that it may be the act of Pol IV transcription that helps clear Pol II from these regions (MCKINLAY *et al.* 2017).

How might the CTD expand the functional repertoire of Pol IV? Our in vitro transcription experiments show that Pol IV assembled using NRPD1-ΔCTD has RNA polymerase activity that is indistinguishable from Pol IV assembled using full-length NRPD1, at least under conditions in which transcription takes place using only an oligonucleotide DNA template and an RNA primer. Moreover, Pol IV interacts with RDR2 regardless of whether full-length NRPD1 or NRPD1-ΔCTD is used to assemble the holoenzyme, such that there is no reason to suspect that an ability to synthesize double-stranded P4R2 RNAs, which are the precursors of siRNAs, should be impaired. We speculate that although the CTD is not needed for the fundamental catalytic activity of the Pol IV-RDR2 complex, the reduced siRNA levels in vivo in the absence of the CTD likely indicates an impairment in Pol IV activity in the context of chromatin. We hypothesize that the CTD may be an important for interactions with chromatin remodelers or other proteins that facilitate Pol IV recruitment, elongation, termination on RNA processing. Lending credence to this hypothesis, the DeCL domain within the CTD of the Pol V largest subunit, NRPE1 is similarly required for production of Pol V transcripts *in vivo* but not *in vitro* (WENDTE *et al.* 2017).

## Materials and Methods

### Plant material

*Arabidopsis thaliana* mutant lines *nrpd1-3* and *nrpe1-11* have been described previously (ONODERA *et al.* 2005; PONTIER *et al.* 2005), as have full length NRPD1-Flag (*nrpd1-3*) and NRPB2-Flag (*nrpb2*) transgenes and *nrpb2* (PONTES *et al.* 2006; ONODERA *et al.* 2008). All plants were grown on soil in long day conditions (16 hours light, 8 hours dark).

### Generation of the nrpd1 ΔCTD transgenic line

The C-terminal domain deletion of NRPD1 was generated by modification of a pENTR-NRPD1 full-length genomic clone that includes the native gene promoter (PONTES *et al.* 2006). Using this clone as the DNA template, PCR primers were used to generate a truncated version of the gene lacking the CTD (Table S9). Pfu Ultra DNA polymerase (Stratagene) was used to amplify the sequences. The PCR products were gel purified and cloned into pENTR-TOPO S/D (Invitrogen) before being recombined into pEarleyGate302 to incorporate a C-terminal FLAG epitope (EARLEY *et al.* 2006). The resulting plasmid was transformed into *Agrobacterium tumefaciens* and used to transform *nrpd1-3* plants using the floral dip method (BECHTOLD AND PELLETIER 1998; CLOUGH AND BENT 1998).

### Protein immunoprecipitation and immunoblot analysis

Proteins were extracted from 4 grams of ∼2.5 week old above-ground plant tissues that were ground to a fine powder in liquid nitrogen. The resulting powder was suspended in 14 ml lysis buffer (50 mM Tris HCl pH 7.6, 150 mM NaCl, 5mM MgCl_2_, 10% glycerol, 0.5 mM DTT, 0.1% IGEPAL, 1% Plant protease inhibitors (Sigma)), filtered through two layers of miracloth and subjected to centrifugation at 16,000 X g for 15 minutes at 4°C. The resulting supernatant was incubated with anti-Flag Agarose (Sigma) for 2 hours at 4°C on a rotating mixer. The agarose resin was then washed twice with lysis buffer and boiled in SDS PAGE buffer. SDS-PAGE was conducted using Tris-glycine gels and proteins were then transferred to nitrocellulose membranes for immunoblotting. Antibodies were diluted in TBST + 5% (w/v) nonfat dried milk as follows: 1:500 anti-NRP(D/E)2, 1:250 anti-RDR2, 1:500 anti-NRP(B/D/E)11, and 1:2,000 anti-Flag-HRP (Sigma). Anti-rabbit-HRP (Santa Cruz Biotechnology) diluted 1:5,000 was used as secondary antibody to bind the primary antibodies. Native antibodies to NRP(D/E)2, RDR2, and NRP(B/D/E)11 were previously described (ONODERA *et al.* 2005; HAAG *et al.* 2009; WENDTE *et al.* 2017).

### Nuclear immunolocalization

Immunolocalization studies were conducted as described previously (Pontes et al., 2006), using nuclei of 4-week old plants fixed in 4% formaldehyde and incubated with antibodies recognizing the C-terminal Flag epitopes fused to the NRPD1 recombinant proteins.

### In vitro transcription assays

*In vitro* transcription was conducted as described in (HAAG *et al.* 2012) as modified in (WENDTE *et al.* 2017) using Pol IV extracted using lysis buffer ((50 mM Tris HCl pH 7.6, 150 mM NaSO_4_, 5mM NaSO_4_, 10% glycerol, 0.5 mM DTT, 1% Plant protease inhibitors (Sigma)) and affinity captured on anti-Flag agarose (Sigma). Pol IV associated resin was washed once with CB100 buffer (100 mM potassium acetate, 25 mM HEPES, pH 7.9, 20% glycerol, 0.1 mM EDTA, 0.5 mM DTT, 1 mM PMSF), then resuspended in a mix of 50 μL CB100 buffer and 50 μL 2x transcription reaction buffer (120 mM ammonium sulfate, 40 mM HEPES, pH 7.6, 20 mM magnesium sulfate, 20 μM zinc sulfate, 20% glycerol, 0.16 U/μL RNaseOUT, 20 mM DTT, 2 mM ATP, 2 mM UTP, 2 mM GTP, 0.08 mM CTP, 0.2 mCi/mL alpha ^32^P-CTP and 4 pmols of template). Transcription reactions were conducted at room temperature for 60 minutes on a rotating mixer, and stopped by addition of 50 mM EDTA and heating at 75°C for five minutes. Transcription products were enriched using PERFORMA spin columns (EdgeBio) and precipitated using 1/10 volume of 3M sodium acetate, pH 5.2, 20 μg glycogen and 2 volumes isopropanol at −20°C overnight. Radioactive RNA transcripts were resolved on 15% denaturing polyacrylamide gels, transferred to Whatman 3MM filter paper, dried under vacuum and visualized by phosphorimaging. Template sequences are found in Table S9.

### Whole genome bisulfite sequencing and analysis

Bisulfite sequencing and analysis was conducted as described in (WENDTE *et al.* 2017). DNA (100 ng) extracted from ∼2.5 week old above ground plant tissues was prepared using the Illumina TruSeq DNA methylation library prep kit according to the manufacturer’s instructions. Libraries were sequenced using an Illumina NextSeq instrument. Base calling, adapter trimming, and read size selection (>≠ 35bp) were performed using bcl2fastq v2.16.0.10. To counter methylation end bias, the first 7 and last 2 bases of each read were removed and a minimum q score of 25 was applied as a filter using Cutadapt version 1.9.1 (MARTIN 2011).

Mapping of sequenced reads to the *Arabidopsis thaliana* TAIR10 genome, removal of PCR duplicates, and extraction of methylation information for cytosines with a minimum coverage of 5 reads was completed using Bismark version 0.16.1 with default settings (KRUEGER AND ANDREWS 2011). The bisulfite conversion rate was calculated based on the number of methylated cytosines divided by total number of mapped cytosines (converted and un-converted) of the chloroplast genome, which is unmethylated.

Differently methylated regions (DMRs) for CHH methylation relative to Col-0 were defined using the R package methylKit version 0.9.5 (AKALIN *et al.* 2012). The genome was analyzed in 300 base pair sliding windows with a step size of 200 base pairs. Windows were counted if 10 informative cytosines with at least 5 reads each were observed. Significant hypo-DMRs, where methylation is redued relative to WT, were assessed at two cutoff values: ≥ 10% decrease or ≥ 25% decrease in methylation. DMRs were calculated using a logistic regression and significance was assumed for q-values less than or equal to 0.01, with p-values corrected for multiple testing using the SLIM method (AKALIN *et al.* 2012). Overlapping DMRs were merged into a single region prior to determining the number of DMRs in each line.

To quantify percent methylation within regions of interest, the methylKit regionCounts function was utilized with the genomic coordinates of interest input as a bed file (AKALIN *et al.* 2012). In box plots, outliers 1.5 times the interquartile range beyond the upper or lower quartile were omitted.

To rank Pol IV-dependent DMRs based on % methylation change in *nrpd1* ΔCTD relative to Col-0 (as shown in Figure 3), only regions that also had a minimum coverage of 25 reads in each Col-0 sRNA sequencing replicate were considered.

To calculate % methylation values for the locus-specific analyses of Figures 4 and 5, % methylation was summed for all Pol IV-dependent DMRs found to overlap the locus of interest, including 300 base pairs of 5’ and 3’ flanking sequence.

Accession numbers for previously published bisulfite sequencing data are provided in Table S1.

### sRNA analysis

sRNA blots were conducted on RNA extracted from 350 mg inflorescence tissue as described previously (WENDTE AND PIKAARD 2017).

Whole genome small RNA sequencing was conducted using 1 μg total RNA isolated from 2.5 week old above-ground plant tissues isolated using Trizol. Libraries were prepared using the Illumina TruSeq small RNA library prep kit according to the manufacturer’s instructions, except that the size selection step was adjusted to select for RNAs of 15-60 nt.

Raw reads were adapter and quality (q > 20) trimmed and size selected (16-60 nucleotides) using Cutadapt version 1.9.1 (MARTIN 2011). Reads were first filtered to remove structural RNAs (tRNAs, rRNAs, snRNAs, and snoRNAs) then mapped to the *Arabidopsis* TAIR10 genome using ShortStack version 3.4 default settings (JOHNSON *et al.* 2016). 24 nt siRNA clusters were defined with a minimum coverage of 25 reads, comprised of at least 80% 24 nt siRNAs. Clusters with 75 base pairs were merged into a single region. Only clusters that were identified in both Col-0 WT replicates were analyzed.

Small RNA read counts for regions of interest were extracted from BAM files using the ShortStack --locifile file function (JOHNSON *et al.* 2016). Counts were normalized as reads per million (RPM) based of total mapped reads. To calculate the log_2_ fold change relative to Col-0, 0.5 reads were first added to the read count for each region of interest to account for 0 values. After read count normalization, the average RPM of wild-type Col-0 was calculated and log_2_ fold changes were calculated for each sample as: log_2_ (Sample RPM / Avg. wild-type Col-0 RPM). In box plots, outliers 1.5 times the interquartile range beyond the upper or lower quartile were omitted. Accession numbers for previously published sRNA data are listed in Table S1.

To analyze Pol IV dependent precursor RNAs, reads size 26-60 nt were summed for each 24 nt siRNA cluster in each Col-0 replicate and only those regions with a minimum coverage of 10 reads for precursor RNA size classes were analyzed for each genotype.

To calculate log_2_ fold change values for the locus-specific analyses in Figures 4 and 5, read counts were summed for all 24nt siRNA clusters found to overlap the locus of interest with 300 base pairs of 5’ and 3’ flanking sequences included.

### Chop PCR

Genomic DNA was digested with *Hae*III alone (Figure 1) or double-digested with *Alu*I and *Hae*III (Figure 3) (NEB) at 37 °C for 3 hours, followed by PCR using GoTaq Green (Promega) with primers that flank enzyme cut sites listed in Table S9. For no-digest controls, DNA was subjected to the same protocol but with no restriction enzymes added.

### RT-PCR

RNA extraction of 2.5 week above-ground tissues was conducted using Trizol. Reverse transcription was conducted using SuperScript III (Invitrogen), 1-5μ RNA, and random primers (Sigma) or dN6 primers (NEB). PCR amplification conducted using GoTaq Green (Promega) with primers listed in Table S9.

### Data and reagent availability

Plant strains are available upon request. Deep sequencing data generated in this study have been deposited in NCBI's Gene Expression Omnibus (EDGAR et al. 2002) and are accessible through GEO Series accession number GSE95825. Whole genome sRNA and bisulfite sequencing statistics and data accessions numbers are provided in Table S1. Genomic coordinates of differentially methylated regions is provided in Table S2. Numerical values for percent CHH methylation at Pol IV-dependent loci is provided in Table S3. Genomic coordinates for siRNA clusters are provided in Table S4. Genomic coordinates and small RNA values for 500 loci whose methylation is most or least affected by deletion of the CTD are provided in Tables S5 and S6. Oligonucleotides used in the study are provided in Table S9.

## Acknowledgements

The authors thank James Ford of the Indiana University Center for Genomics and Bioinformatics for help with library preparation and sequencing. This work was supported by funds to C.S.P. as an Investigator of the Howard Hughes Medical Institute and Gordon and Betty Moore Foundation and from grant GM077590 from the National Institutes of Health. J.M.W. was supported by NIH training Grant, T32GM007757, and predoctoral fellowship Award F31GM116346. The content of this work is solely the responsibility of the authors and does not necessarily represent the views of our sponsors. The authors declare no conflicts of interests. J.M.W. and J.R.H. designed and performed experiments, analyzed data, and wrote the manuscript. J.S. performed *in vitro* assays. O.P. performed nuclear localization. S.M. conducted Chop-PCR assays on the *nrpd1* ΔCTD / *nrpe1-11* double mutant. C.S.P was involved in designined experiments, analyzing data, and writing the manuscript.

